# Splicing-dependent transcriptional activation

**DOI:** 10.1101/2022.09.16.508316

**Authors:** Maritere Uriostegui-Arcos, Steven T. Mick, Zhuo Shi, Rufuto Rahman, Ana Fiszbein

## Abstract

Transcription and splicing are intrinsically coupled. Transcription dynamics regulate splicing, and splicing feeds back to transcription initiation to jointly determine gene expression profiles. A recently described phenomenon called exon-mediated activation of transcription starts (EMATS) shows that splicing of internal exons can regulate transcription initiation and activate cryptic promoters. Here, we present the first complete catalog of human EMATS genes that have a weak alternative promoter located upstream and proximate to an efficiently spliced internal skipped exon. We found that EMATS genes are associated with Mendelian genetic diseases —specifically intellectual development disorders, cardiomyopathy, and immunodeficiency— and provide a list of EMATS genes with pathological variants. EMATS was originally described as a natural mechanism used during evolution to fine-tune gene expression through punctual genomic mutations that affect splicing. Here, we show that EMATS can be used to manipulate gene expression with therapeutic purposes. We constructed stable cell lines expressing a splicing reporter based on the alternative splicing of exon 7 of SMN2 gene under the regulation of different promoters. Using a small molecule (Risdiplam) and an antisense oligonucleotide (ASO) modeled after Spinraza, we promoted the inclusion of SMN2 exon 7 which triggered an increase in gene expression up to 40-folds by activating transcription initiation. We observed the strongest effects in reporters under the regulation of weak human promoters, where the highest drug doses dramatically increased exon inclusion. Overall, our findings present evidence to develop the first therapeutic strategy to use EMATS to activate gene expression using small molecules and ASOs that affect splicing.

## Introduction

RNA splicing is a highly regulated process that influences almost every aspect of eukaryotic cell biology. Early studies suggest that both spliceosome assembly and catalysis of splicing occur in a co-transcriptional manner (Custódio and Carmo-Fonseca, 2016; Oesterreich et al., 2016; Osheim et al., 1985), creating opportunities for functional connections between transcription and splicing. A key player mediating the effects of transcription on RNA splicing is the RNA polymerase II (RNAPII) (Zhang et al., 2021) itself as post-translational modifications of its C-terminal domain (CTD) can create a binding platform for splicing factors (recruitment model), as well as affect the rate of transcription elongation that modulates downstream splicing decisions (kinetic model) (Fong et al., 2014; Kornblihtt et al., 2013; de la Mata et al., 2003). Since chromatin structure can influence transcription rate and recruitment of factors, histone modifications are a powerful source of splicing regulation (Fiszbein and Kornblihtt, 2016; Fiszbein et al., 2016). More recently, accumulating evidence suggests a “reverse-coupling” mechanism in which splicing feeds back to transcription (Furger et al., 2002; Shaul, 2017). Adding an intron to an intron-less gene often boosts different stages of gene expression in plants, fungi, and animals by a phenomenon known as intron-mediated enhancement of gene expression (Shaul, 2017). Moreover, a functional 5’-splice site in close proximity to a promoter was found to enhance transcriptional output (Furger et al., 2002), and change chromatin structure (Bieberstein et al., 2012). We recently characterized a related phenomenon called Exon-Mediated Activation of Transcription Starts (EMATS), in which the splicing of internal exons impacts the spectrum of promoters used and the expression levels of the host genes (Fiszbein et al., 2019). We observed that EMATS modulates gene expression through changes in splicing efficiency of internal exons during evolution. The strongest effects are seen when there is a highly included skipped exon (SE) located downstream and proximate to a weak promoter. We also showed that specific proteins involved in splicing, whose depletion has large effects on alternative promoter use, have widespread interactions with core transcription machinery and that the splicing factor HNRNPU recruits core transcription machinery locally. Overall, our observations support a reversed “recruitment model” in which specific splicing factors recruit core transcription machinery to the vicinity of transcripts as they are being transcribed, boosting RNAPII occupancy and activity of nearby promoters.

Dysregulations in splicing, spliceosome complexes, and RNA processing can lead to many diseases including spinal muscular atrophy, tauopathies, Hutchinson-Gilford progeria syndrome (HGPS), and hypercholesterolemia. These dysregulated processed are also implicated in several cancers types and especially those with mutations that affect RNA splicing in tumor suppressor genes such as BRCA1, APC, and MLH 1 (Anczuków and Krainer, 2016; Tazi et al., 2009). Thus, a deep dive into understanding how RNA splicing contributes to gene expression programs is critical to uncovering pathophysiological mechanisms of diseases. In EMATS, splicing changes are associated with promoter choice and gene expression levels. These changes can regulate global gene expression networks in a tissue- and specie-specific fashion. While this functional characterization of EMATS was described, the role of EMATS genes in human diseases is still not characterized.

Small molecules that target specific DNA sequences have been widely used to control gene expression (Gottesfeld et al., 1997), and small-molecule drugs that target RNA have also emerged as a new opportunity to therapeutically modulate numerous cellular processes (Warner et al., 2018). More recently, antisense oligonucleotides (ASOs) have emerged as a specific, rapid and potentially high-throughput approach for modulating gene expression through recognition of cellular RNAs (Di Fusco et al., 2019). A subset class of ASOs, the splice-switching oligonucleotides (SSOs), are short, antisense oligonucleotides that base-pair with a pre-mRNA and disrupt the normal splicing pattern by blocking spliceosome or splicing factor recruitment (Roberts et al., 2020). SSOs offer an effective way to alter splicing and have been successfully used to treat Duchenne muscular dystrophy and spinal muscular atrophy (Hua and Krainer, 2017; Kole and Leppert, 2012; Krainer, 2018; Zhu et al., 2016). Recent studies show that chromatin modifiers, such as histone deacetylase inhibitors cooperate with ASOs/SSOs to control splicing (Marasco et al., 2022). However, most of these therapeutics have focused on gene silencing while successful strategies to activate gene expression remain to be developed.

Here, we present a complete catalog of EMATS genes and investigate their role in human genetic diseases. We found several genetic mutations associated with Mendelian diseases that switch isoform expression by EMATS and characterized their molecular functions. Since EMATS can boost gene expression, we developed the first therapeutic strategy to treat associated genetic diseases based on small molecules or ASOs that activate transcription through splicing in EMATS genes.

## Results

### A catalog of human EMATS genes

Working with mouse evolutionarily new exons, we previously observed that EMATS requires splicing of an internal exon, not merely presence of a 5’ or 3’ splice site, to activate a cryptic promoter, and the effect is stronger when the promoter is intrinsically weak and exon splicing is efficient (Fiszbein et al., 2019). EMATS is sensitive to genomic distance between exon and upstream promoter and shows the strongest effects when there are 1-5 kilobases (kb) in between them. We also observed increased ribosome occupancy in isoforms that are activated through EMATS, suggesting that the upregulation of transcription can be magnified by an increase in translation efficiency. Since these changes in gene expression have potential therapeutic benefits, here we aimed to identify human genes that have an EMATS structure in which their transcription and translation activity could be modulated through changes in splicing.

Using the GENCODE (Frankish et al., 2021) annotation we started with almost 20,000 protein coding genes (Fig. 1A). Based on the genome annotations, we collapsed all overlapping exons from the same gene into one “metaexon” that accounts for alternative splice sites, alternative transcription start sites, and alternative polyadenylation sites. We then used the Hybrid-Internal-Terminal (HIT index) method to classify exons in their isoform-specific transcript usage. Exons were determined to be alternative or constitutive by examining substitute exon inclusions in different transcripts. Percent splice-in (PSI) levels of inclusion for alternative exons were measured using the HIT index (Fiszbein et al., 2022) for first and last exons, and using rMATS (Shen et al., 2014) for internal exons across 10 tissues and several individuals using a curated list of samples from the GTEx project. We defined weak alternative exons those with median PSI values across tissues below the median levels of all exons in their same classification, and strongly included alternative exons those with median PSI values above the population median value. We identified human protein coding genes with SEs multiple first exons and selected among those, genes in which the start coordinate of a weak alternative first exon (AFE) is located within 5 kb upstream from the start coordinate of one or more highly included SE (Fig. 1A). Our complete list of 1,463 human EMATS genes include 4,117 unique AFE/SE pairs (Supplementary table 1, Supplementary table 2). A detailed code to identify EMATS genes is provided at https://github.com/fiszbein-lab/emats-genes.

**Figure 1.**
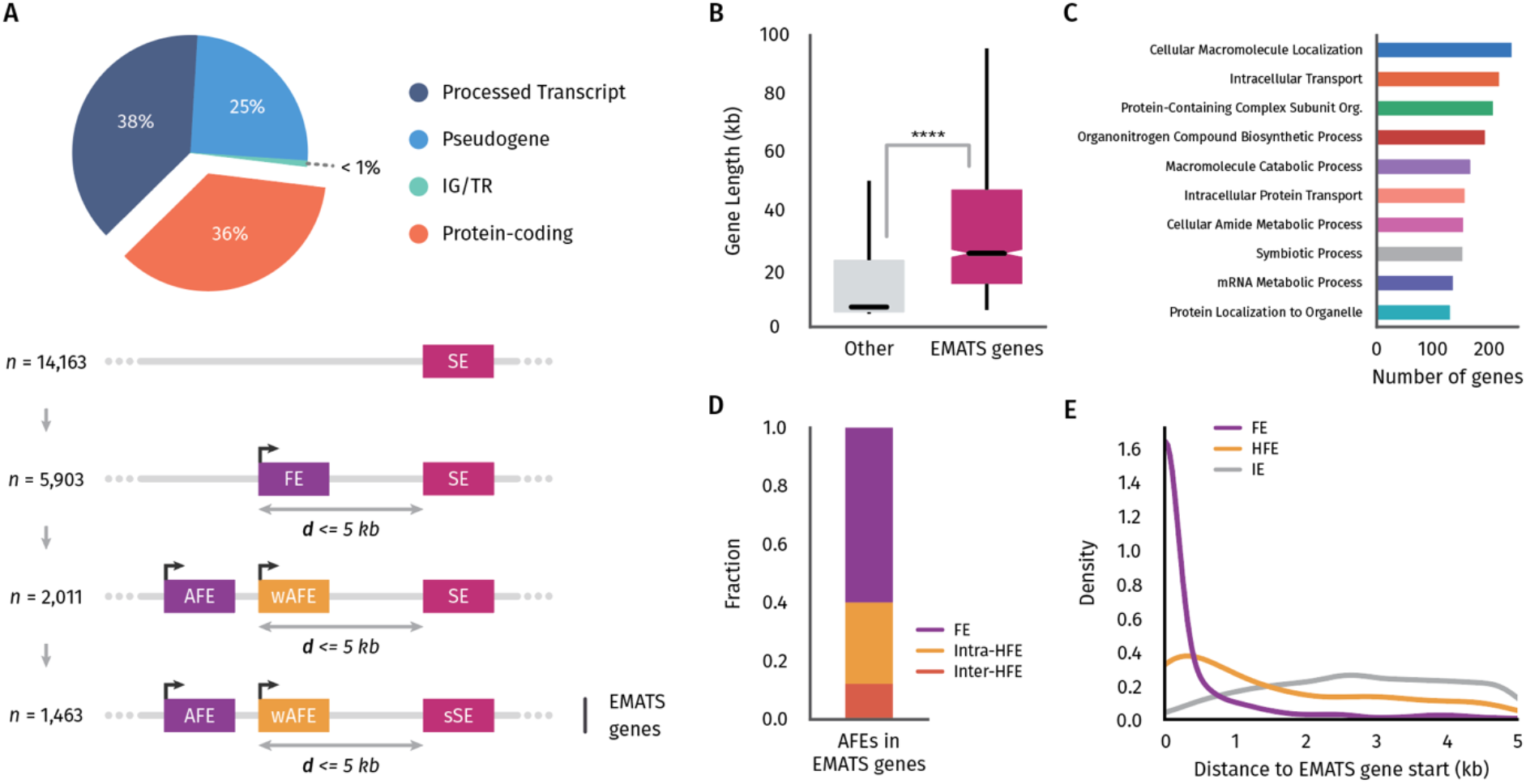
Identification and characterization of human EMATS genes. **A**, a step-by-step identification of human EMATS genes with the following characteristics: protein coding genes, with multiple alternative promoters, and at least one weak AFE located within 5kb upstream of a strong SE. **B**, distribution of total gene length for genes with EMATS structure and genes without EMATS structure. **C**, enrichment of human EMATS genes in gene ontology classifications. **D**, percentage of first exons in an EMATS structure classified as obligated first exons (FE), first internal inter-tissue hybrid exons (inter-tissue HFE), or first internal intra-tissue hybrid exons (intra-tissue HFE). E, relative position of exons including Internal exons (IE) in an EMATS structure within their gene length.

We found that human genes with EMATS structure are longer genes (Fig. 1B) with longer transcripts (Fig. S1A) that present significantly more alternative isoforms compared to non-EMATS genes (Fig. S1B). Consistent with similar analyses in other species, human EMATS genes are enriched in intracellular transport, protein localization, and mRNA metabolism (Fig. 1C). We recently classified a new category of exons that we called hybrid exons which are used as terminal and internal exons in different transcripts (Fiszbein et al., 2022). We identified ∼100,000 human inter-tissue hybrid exons which are used as terminal exons in one tissue but as internal exons in other tissues and ∼20,000 intra-tissue hybrid exons which are used as hybrid within the same tissues. Notably, we found that the majority of AFEs in an EMATS structure are obligate first exons (∼60%), followed by intra-tissue hybrid first internal exons (∼28%), and inter-tissue hybrid first internal exons (∼12%) (Fig. 1D). As expected, obligate first exons in EMATS genes are mostly located towards the 5’ end of genes, while hybrid first internal exons and internal exons in an EMATS structure are distributed more equally within gene bodies towards the 3’ end (Fig. 1E). These findings reinforce the notion that EMATS is a mechanism that influences transcription starts with potential crucial implications in gene regulation without altering protein coding structures.

### EMATS genes are associated with genetic diseases

In order to provide a framework to study the role of EMATS in human diseases, we assembled a master gene-phenotype dataset from Online Mendelian Inheritance in Man (https://omim.org/) to obtain a full list of genes involved in human genetic diseases (Scott et al., 2006). OMIM is a continuously updated compendium of human genes, diseases, and traits with in-depth information on the molecular mechanisms and phenotypic expression. This dataset was merged with the EMATS structure dataset to obtain a list of EMATS genes with connections to human diseases (Fig. 2A). To find variants within these genes that commonly cause disease, we pulled variant data from ClinVar (Landrum et al., 2014, 2020). Variants with pathological or pathological/likely pathological annotations were selected and combined with the above data to identify variants specifically within the exons in EMATS structure. This led to a list of 373 EMATS genes with 3,493 pathological variants affecting either the AFE or the SE located in an EMATS structure (Fig. 2A). We identified genetic diseases involving exons with EMATS structure and found enrichment for intellectual development disorders, including emotional and behavioral disorders, cardiomyopathy, immunodeficiency, myopathy, neurodevelopment disorders, and others (Fig. 2A). We then explored the specific genomic position of the genetic variants and divided them into upstream of EMATS AFE/SE, in TSS/3’splice site, within EMATS AFE/SE, in 5’ splice site, and downstream of EMATS AFE/SE (Fig. 2B). Besides the majority of variations located within EMATS AFE/SEs, a significant number of genetic variations associated with diseases are located near the EMATS TSS in the case of AFEs and the 3’ splice site in the case of SE. We then sought to explore further the consequences of genetic mutations associated with splicing in EMATS genes. We selected splice site mutations of EMATS exons in genes involved in Mendelian diseases identified with OMIM and ClinVar data. EMATS FHL1 gene expresses three different main isoforms by a combination of alternative TSSs and alternatively spliced exons named FHL1A, FHL1B, and FHL1C, all of which are involved in skeletal and cardiac muscle function(Gueneau et al., 2009). More than 50 FHL1 gene mutations have been associated with myopathies. Here, we focus on two mutations that affect splicing of exon 6 and trigger a downregulation of the second TSS by EMATS (Fig. 2C). Patients with these genetic variants express a wild type FHL1C, a mutated version of FHL1B which disrupts a highly conserved double zinc finger domain called LIM domain (Wei and Zhang, 2020), and reduced levels of a mutated version of FHL1A by EMATS.

**Figure 2.**
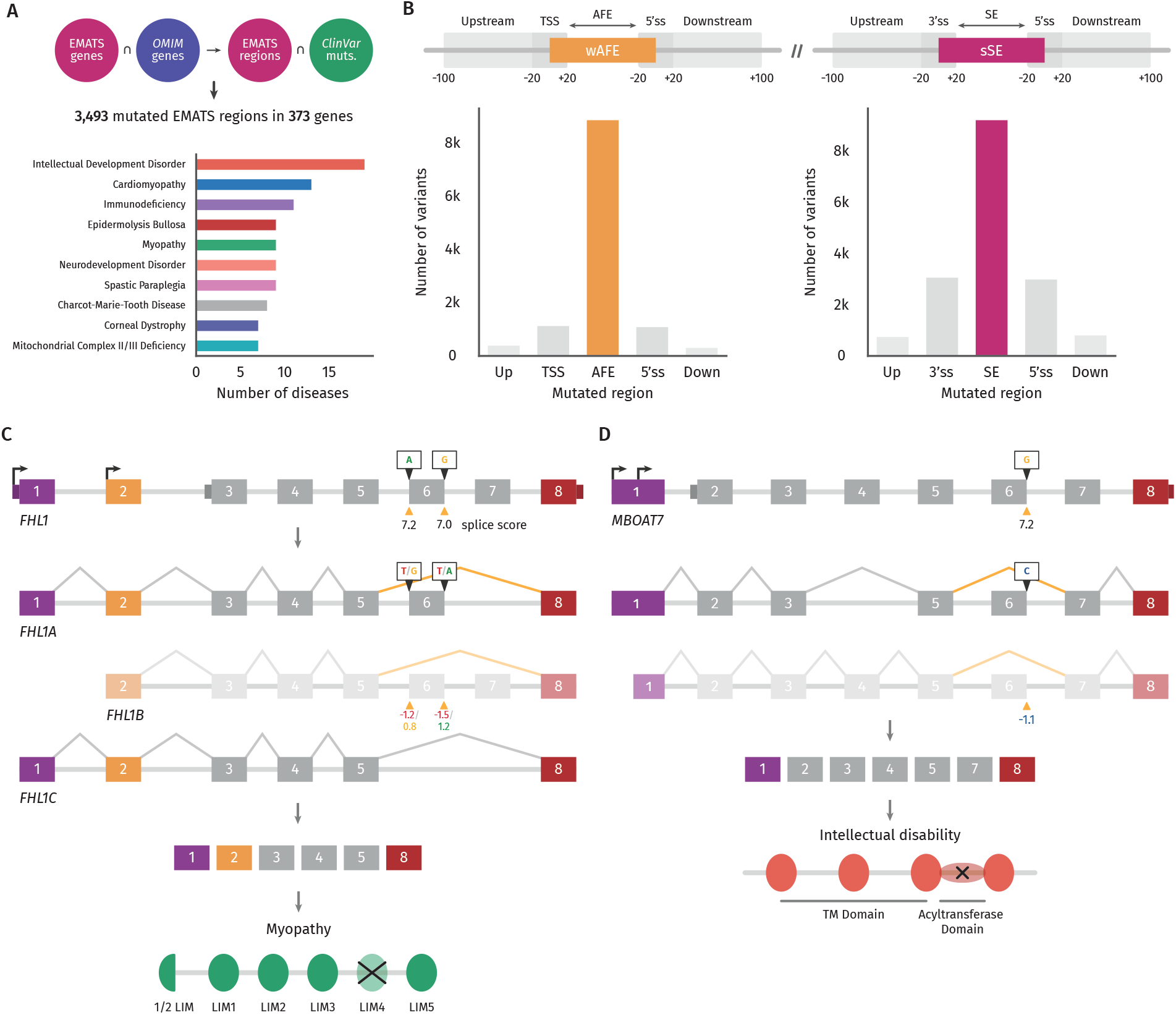
Pathological variants affecting human EMATS genes. **A,** a step-by-step identification of Mendelian pathological genomic variants associated with EMATS regions of human genes with EMATS structure (upper) and Gene Ontology enrichment of genetic human diseases associated with genes with EMATS structure (lower). **B,** number of pathological genomic variants in human genes with EMATS structure located in specific regions of EMATS’ exons. **C, D,** genomic structure of FHL1 (**C**) and MBOAT7 (**D**) genes and their isoforms, location of pathological genomic variants and their effect on splicing efficiency measure by changes in splice score, and disruption of protein domains by EMATS in patients with pathological variants.

These genetic variations that affect splicing and trigger a downregulation of specific isoforms by EMATS are associated with severe muscular dystrophy-like disorders with childhood onsets. In the case of MBOAT7 gene, we found a mutation in the 5’ splice site of exon 6 which decrease the splicing score of that exon to a negative value (Fig. 2D). MBOAT7 encodes a member of the membrane-bound-O-acyltransferases family of integral membrane proteins that have acyltransferase activity. The exclusion of exon 6 would mostly affect the isoform that starts in the downstream TSS by EMATS, favoring expression from the upstream TSS. The predominant isoform codes for a protein with a disruptive acyltransferase domain that is associated with intellectual disability(Johansen et al., 2016) (Fig. 2D). Together, these examples illustrate how EMATS can trigger genetic diseases and magnified disorders triggered by splicing variants.

### Splicing of internal exons is associated with increased gene expression

We recently showed that during evolution, the gain of new exons is associated with increased gene expression levels. Here, we wondered whether the splicing of skipped exons remains associated with host gene expression in human transcriptomes. We first analyzed the association between gene expression and alternative splicing events across 10 human tissues using data from the GTEx project (GTEx Consortium, 2013). Although all splicing events are positively associated with gene expression, splicing of skipped exons showed the largest effect (Fig. 3A). Consistent with our findings during evolution, higher inclusion levels of skipped exons measured in SE percent splice in (PSI) are associated with higher gene expression levels across human tissues in a global scale (Fig. 3B) and for individual genes (Fig. 3C). We featured four individual genes with the highest associations between inclusion levels of SE and gene expression across human tissues and samples. In these cases, the internal SE was located between 3 and 11 kb downstream of the adjacent promoter, suggesting that the location of the SE and the TSS also plays a role in their regulation. Notably, the association between splicing of skipped exons and gene expression levels is stronger in EMATS genes (Fig. 3D). This observation opens up the possibility of new therapeutic strategies by manipulating gene expression of genes associated with human genetic diseases through changes in splicing.

**Figure 3.**
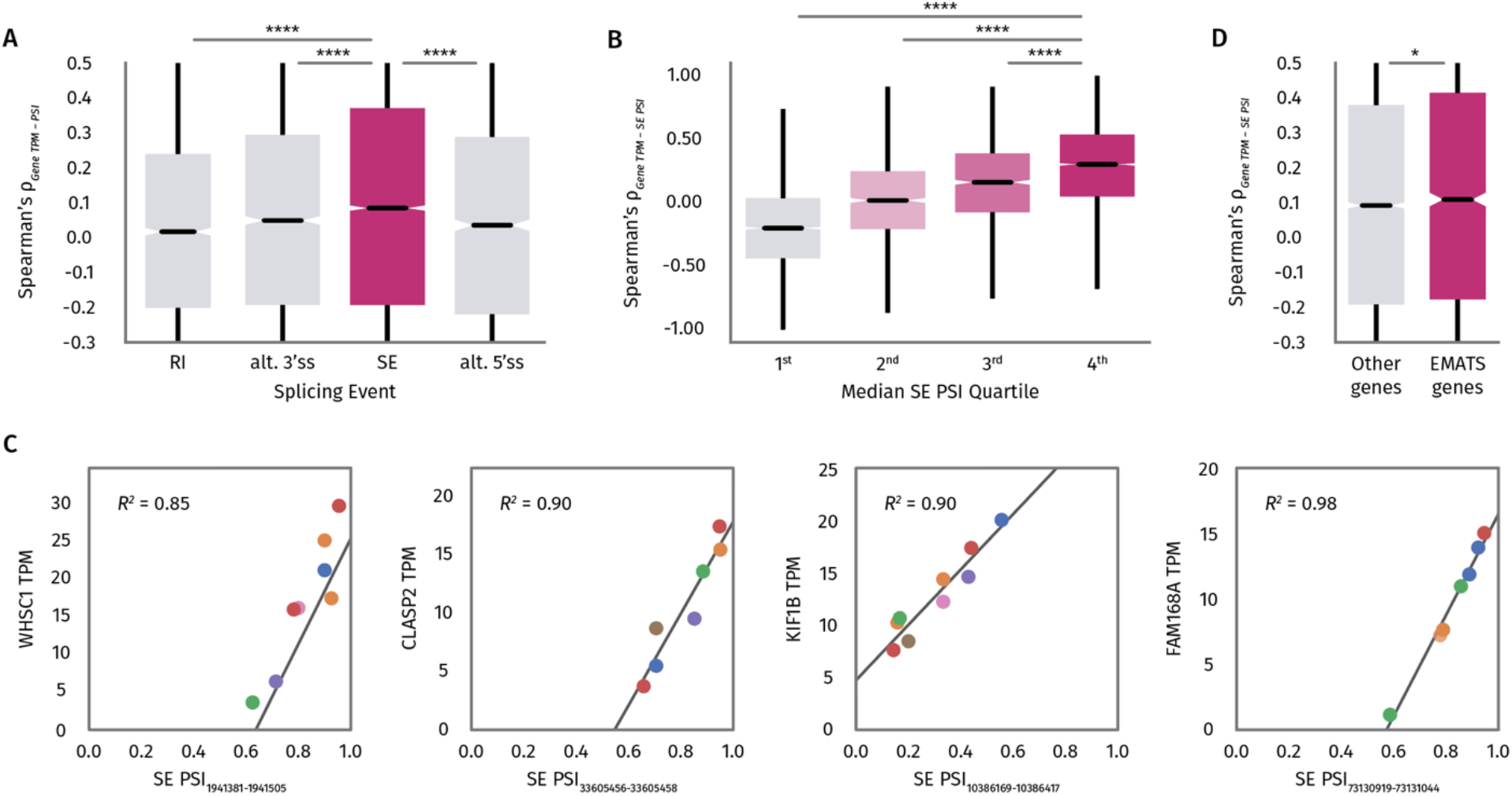
Splicing efficiency and increased gene expression are coupled. **A,** distribution of the spearman correlation between gene expression in TPMs and splicing events (intron retention, alternative 3’ splice sites, skipped exons, and alternative 5’ splice sites) in PSI values across human tissues and samples. **B,** distribution of spearman correlation between gene expression and SE PSI across human tissues and samples binned in quartiles of SE PSI. **C,** linear regression of gene expression versus SE PSI for individual genes with the highest associations. **D,** distribution of the spearman correlation between gene expression and SE PSI across human tissues and samples for non-EMATS genes compared to EMATS genes.

In order to analyze the associated changes in splicing ratios and gene expression during cellular transitions and considering that EMATS genes are enriched for immunological disorders, we then calculated the correlation between changes in splicing of skipped exons with changes in gene expression during SARS Cov-2 infection. We observed that EMATS genes show a significantly stronger association between splicing of skipped exons and gene expression levels during viral infection compared to non-EMATS genes (Fig. S2A). Consistent with our previous findings, the effects are stronger when there is an AFE located proximal and upstream of the SE (Fig. S2B). Altogether, these observations indicate that splicing of SEs is associated with gene expression in human transcriptomes and their regulation is positively correlated during cellular transitions, with the strongest effects in EMATS genes.

### Splicing boost gene expression

Since splicing of internal exons is associated with gene expression in humans, we wondered whether splicing perturbations can be used to control gene expression. Moreover, since the association between splicing and gene expression is stronger in EMATS genes, we explored the possibility of using EMATS to develop a therapeutic strategy to treat genetic diseases based on splicing modulation. We already showed that inhibition of splicing with ASOs can down-regulate expression from the host genes through EMATS (Fiszbein et al., 2019). Here, we investigate whether splicing activation can be used to boost gene expression through EMATS as a potential therapeutic strategy.

First, we generated stable cell lines expressing a splicing reporter based on the well-characterized SMN2 locus associated with muscular dystrophy (Fig. 4A) and triggered splicing changes with small molecules to evaluate potential modulations in gene expression levels (Hua and Krainer, 2017). Treatment with Risdiplam (Markati et al., 2022; Poirier et al., 2018), a small molecule designed to upregulate splicing of SMN2 alternative exon 7, increased inclusion of the alternative exon in our system by several folds (Fig. 4B). The splicing effect correlated with drug concentration upregulating splicing in up to 100-fold with the highest concentration. Notably, the upregulation of splicing was significantly associated with an increase in gene expression of the SMN2 reporter (Fig. 4C). An increase of 100-fold in splicing was associated with a ∼45-fold increase in gene expression. The effect followed a linear pattern with bigger effects on splicing associated with bigger effects on gene expression (Fig. 4D). These observations demonstrate that small molecules that increase inclusion of alternative exons can activate gene expression.

**Figure 4.**
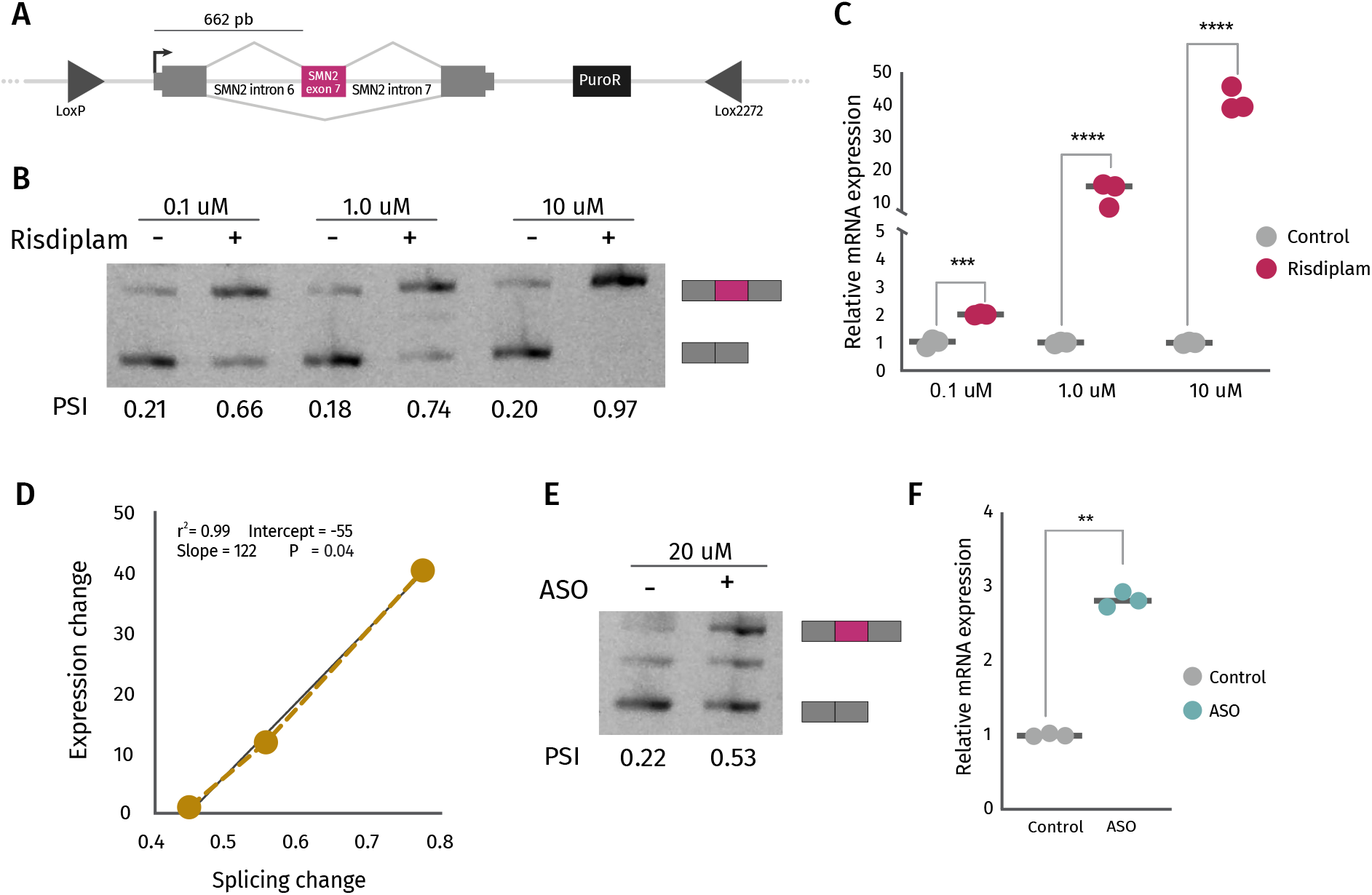
Small molecules and ASOs increase SMN2 exon 7 inclusion and boost gene expression. **A,** diagram of SMN2 splicing reporter in the context of the integration plasmid used to generate HEK293T-A2 stable cells expressing SMN2 splicing reporter under the regulation of the CMV promoter. **B,** stable HEK293T-A2 cells expressing SMN2 splicing reporter were treated with Ethanol (-) or 0.1 μM, 1.0 μM, and 10 μM, of Risdiplam for 24 h. Inclusion of alternative exon 7 in SMN2 was evaluated by RT-PCR. Quantification of densitometric analyses was carried out with Fiji, +Exon/- Exon ratios are shown at the bottom of each lane. **C,** SMN2 expression was evaluated by RT-qPCR after Risdiplam treatment. Results from three independent experiments are shown. Significance was calculated using the student *t* test: ****p<0.0001, ***p<0.001. **D,** linear regression of splicing changes (quantification of B) and gene expression changes (quantification of C) of SMN2 splicing reporter after Risdiplam treatment. **E,** stable HEK293T-A2 cells expressing SMN2 splicing reporter were treated with a scramble antisense oligonucleotide (-) or a specific ASO modeled after the Spinraza (+) for 24 h. Inclusion of alternative exon 7 in SMN2 was evaluated by RT-PCR. Quantification of densitometric analyses was carried out with Fiji, +Exon/-Exon ratios are shown at the bottom of each lane. **F,** SMN2 expression was evaluated by RT-qPCR after ASO treatment. Results from three independent experiments are shown. Significance was calculated using the student t-test: **p<0.01.

We next sought to explore whether ASO targeting splicing have similar effects as small molecules. We used an ASO modeled after the Spinraza (Krainer, 2018) that is known to upregulate splicing of SMN2 exon 7 by blocking an intronic splicing silencer located downstream of exon 7 (Singh and Singh, 2018). Like what we observed with small molecules, the Spinraza-like ASO was able to upregulate splicing of SMN2 under the regulation of the cytomegalovirus (CMV) promoter (Fig. 4E) and increase gene expression levels up to 3-fold (Fig. 4F). This observation is consistent with the original study of SMN2 exon 7 targeting ASOs in which ASOs that promoted exon 7 inclusion of the endogenous locus increased full-length SMN protein levels (Hua et al., 2007). Overall, our findings demonstrate that small molecules and ASOs targeting splicing are an effective method to activate gene expression.

### Splicing can activate transcription from different promoters

In order to investigate whether the upregulation of gene expression by small molecules and ASOs that target splicing could be generalized to other promoter sequences, we performed similar experiments using the splicing reporter under the regulation of different promoter sequences. For these experiments, we used three natural human promoter sequences of different strength (alpha-globin, Fibronectin, and KPTN) and include two mutant versions (a minimal CMV promoter containing only 39bp, and a mutant of the Fibronectin natural promoter in which the CRE at position -170 and the CCAAT box at position -150 have been disrupted by introducing point mutations that abolish binding of the corresponding transcription factors (Cramer et al., 1999)). We observed an increase in inclusion of the alternative exon with Risdiplam treatment under the regulation of all natural promoter sequences and mutants tested (Fig. 5A). Like the effect under the CMV promoter, the splicing regulation increased with higher drug concentrations for all promoters tested. In all cases, the upregulation of SMN2 exon 7 inclusion was associated with an increase in gene expression levels of our reporter (Fig. 5B). These findings indicate that upregulation of splicing of internal exons with small molecules can be used to activate gene expression levels independently of the promoter used.

**Figure 5.**
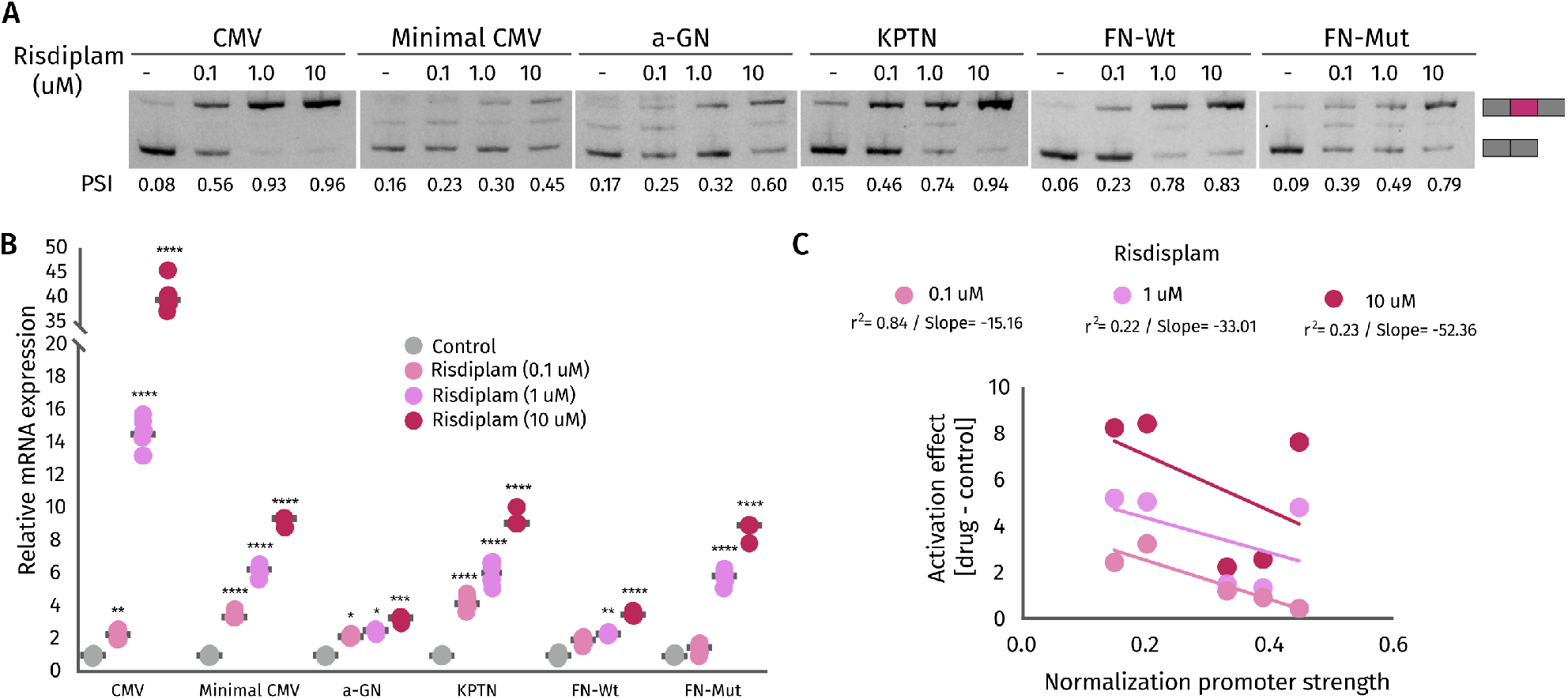
Larger effects of splicing on gene expression with efficient exon inclusion and weak promoters. **A,** HEK293T-A2 stable cell lines expressing SMN2 splicing reporter under the regulation of six different promoters (CMV, Minimal CMV, alfa-globin (a-GN), KTPN, Fibronectin wild type (FN-Wt), and Fibronectin mutated (FN-Mut)) were treated with Ethanol (-) or 0.1 μM, 1.0 μM, and 10 μM of Risdiplam for 24 h. Inclusion of alternative exon 7 in SMN2 was evaluated by RT-PCR for all stable cell lines under the regulation of the different promoters. Quantification of densitometric analyses was carried out with Fiji, +Exon/-Exon ratios are shown at the bottom of each lane. **B,** SMN2 expression was evaluated by RT-qPCR after Risdiplam treatment in all stable lines. Results from three independent experiments are shown. Significance was calculated using Two-way ANOVA with corrections for multiple comparisons, comparing each treatment mean with each control per promoter. \*\*\*\**p*<0.0001, \*\*\**p*<0.001, **p<0.01, *p< 0.05. **C,** linear regression of gene expression changes in SMN2 splicing reporter following Risdiplam treatment with different doses for stable lines under the regulation of the four different human promoters tested and promoter strength measured by basal RT-qPCR in control conditions.

As discussed previously, the EMATS effect is stronger when the splicing of the alternative exon is efficient, and the proximal promoter is weak. To analyze whether the effect of small drugs in gene expression through splicing follow the same or different rules, we quantified the drug-induced effect on gene expression for all different promoters tested and all different drug concentrations. We found that the effect of splicing in gene expression is stronger for higher drug concentrations which are associated with higher splicing efficiency (Fig. S3A-E), suggesting that efficient splicing is associated with a larger effect in gene expression. Moreover, we found that the effect of small drugs in gene expression followed a linear pattern with larger effects associated with weaker promoters (Fig. 5C). Thus, consistent with our previous analyses, splicing effects on transcription are stronger when splicing is efficient, and promoters are weak.

We also tested whether the gene expression activation through splicing with ASOs was also generalized to different promoter sequences. Like the effect with small drugs, ASOs that activate splicing also boost gene expression for all promoter sequences tested (Fig. 6A-B) with higher effects observed for weak promoters (Fig. S4A). Finally, to test what level of gene expression was altered by splicing changes induced with small drugs, we analyzed newly synthesized RNA levels of our splicing reporter following splicing activation with the highest drug concentration. We observed a significant effect of splicing in nascent RNA levels of the reporter (Fig. 6C). Notably, the magnitude of the effect varied with promoter sequences. The effect in nascent RNA under the regulation of a strong CMV promoter was 25% of the effect in total gene expression with the same promoter, while the nascent RNA effect for a weaker alpha-globin promoter was 167% of the gene expression effect (Fig. 6C). These findings suggest that while the effect of splicing in gene expression induced by small drugs is at the transcriptional level, downstream steps of gene regulation may also play a role, and the magnitude of that role depends on the promoter sequence. As noted for steady-state RNA, small molecules that activate splicing also boost transcription initiation with the highest effects observed for weak promoters (Fig. S4B).

**Figure 6.**
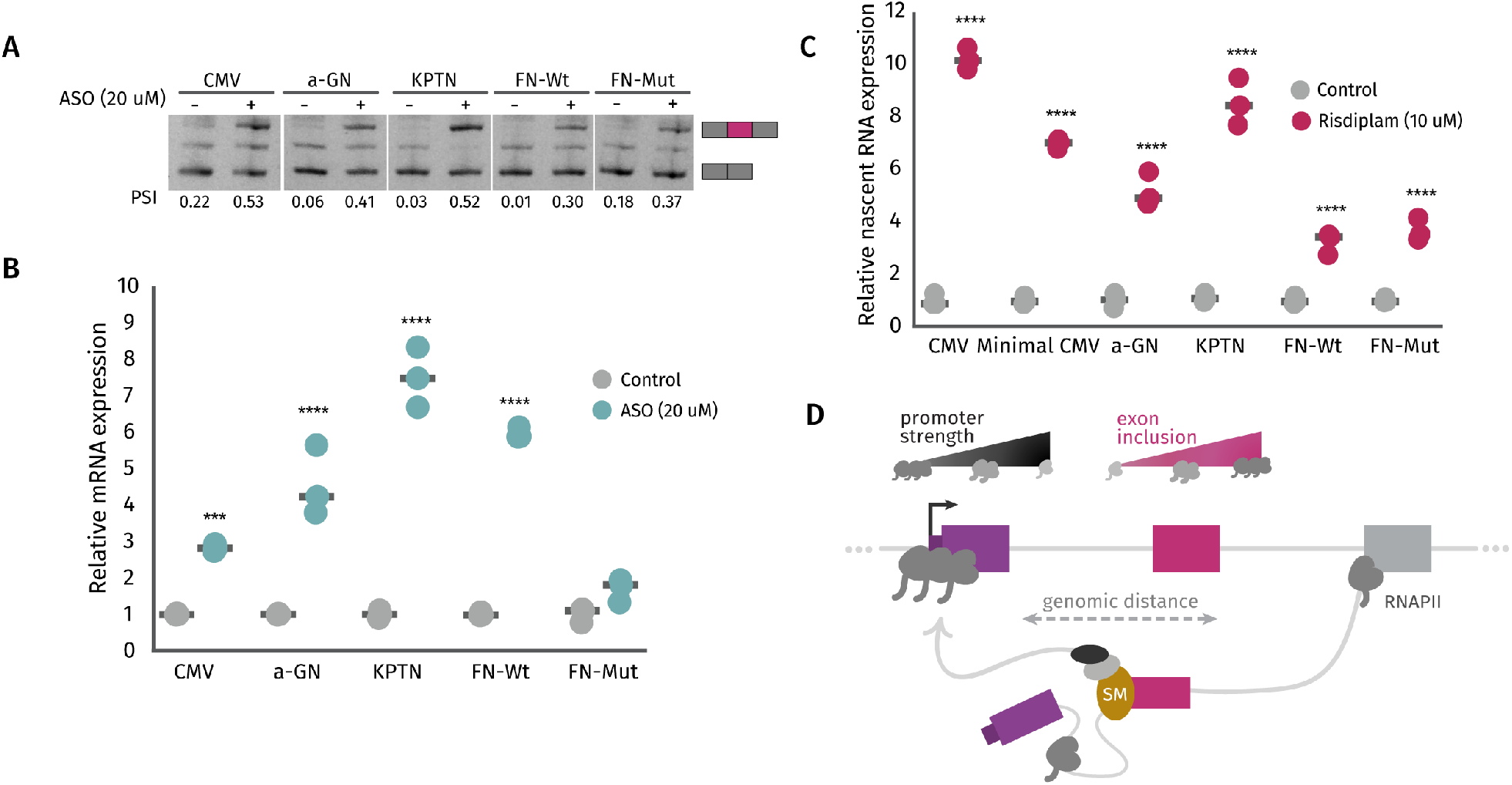
EMATS act at the nascent RNA level. **A, B,** HEK293T-A2 stable cell lines expressing SMN2 splicing reporter under the regulation of five different promoters (CMV, alfa-globin (a-GN), KTPN, Fibronectin wild type (FN-Wt) and Fibronectin mutated (FN-Mut)) were treated with a scramble antisense oligonucleotide (-) or a specific ASO modeled after the Spinraza (+) for 24 h. Inclusion of alternative exon 7 in SMN2 was evaluated by RT-PCR and total gene expression by RT-qPCR. **C,** 30-min nascent RNA was extracted from HEK293T-A2 stable cell lines under the regulation of six different promoters after Risdiplam treatment. SMN2 expression was evaluated by RT-qPCR. Significance was calculated using Two-way ANOVA with corrections for multiple comparisons, comparing each treatment mean with each control per promoter. \*\*\*\**p*<0.0001, \*\*\**p*<0.001, **p<0.01, *p<0.05. **D,** model showing that splicing can activate transcription initiation depending on three main factors: the strength of the promoter, the inclusion of the skipped exon, and the genomic distance between them.

Altogether, our findings demonstrate that small drugs and ASOs that activate splicing can be used to boost gene expression at the transcriptional level from different human promoters, showing the largest effects with higher splicing efficiencies and weaker promoters.

## Discussion

Here, we provide the first comprehensive list of human EMATS genes and establish their link to Mendelian diseases. EMATS genes are defined as protein-coding genes with highly included skipped exons located less than 5 kb downstream of a weak alternative promoter (Fiszbein et al., 2019). Working with this definition, here we identify hundreds of genetic human variants that disrupt splicing of SEs in EMATS genes. Since the splicing of a SE can control the usage of alternative promoters in EMATS genes, we predict that the effect of those genetic variants in splicing is magnified by a secondary effect of splicing perturbation in promoter usage and gene expression levels. Downregulation of specific mRNAs by EMATS have implications in gene expression, and most importantly in the pool of protein isoforms that are expressed from a single gene. As an example, we provided evidence for protein structure disruption in specific EMATS genes caused by genetic variants.

Connections between splicing and gene expression regulation across tissues and species have been observed. Our findings support further broadening of this connection suggesting a direct control of gene expression through splicing of internal exons specifically positioned. In a previous study, we have shown that the mechanism behind the exon-mediated gene expression control might be associated with direct recruitment of transcription machinery to nearby upstream promoters through splicing factors (Fiszbein et al., 2019). Here we reinforced this hypothesis by showing a direct effect of small molecules and ASOs that target splicing in the control of transcription initiation of a splicing reporter single integrated in human cells.

We previously have shown that in EMATS genes the splicing of the skipped exon can control the usage of alternative promoters and regulate the expression of the host gene. Specifically, we have shown that splicing inhibition of a skipped exon in an EMATS gene by splice site-targeting mutations or antisense morpholinos can down-regulate gene expression of the EMATS host gene by several folds. Over the past decade, several techniques have been developed to downregulate the expression of therapeutic-relevant genes. The most characterized technique is the post-transcriptional gene silencing mechanism known as RNA interference (RNAi) triggered by double-stranded RNA (dsRNA), which induces the formation of a ribonucleoprotein complex that mediates sequence-specific cleavage of the transcript cognate with the input dsRNA (Fire et al., 1998). Less common down-regulation techniques include the usage of artificial microRNAs, recombinant nucleic acid molecules with hair pin structures, cationic polymers and single-stranded ribonucleotide oligomers (Roberts et al., 2020), and U1snRNP adaptors (Goraczniak et al., 2009). However, only a few strategies have been proposed to upregulate gene expression with therapeutic benefits. Since in most of the diseases associated with EMATS genes studied here, and several non-Mendelian diseases, gene upregulation would provide a more powerful strategy than gene downregulation, we tested our system as a therapeutic alternative to increase gene expression through splicing activation. Using a small molecule that dramatically increases inclusion of SMN2 exon 7, we were able to upregulate gene expression of the host gene up to 45-fold. Nascent RNA analyses showed that this effect is partially explained by an increase in transcription initiation and suggest other steps of gene regulation at the RNA level are affected. Since previous studies indicate that the effect of splicing in transcription is magnified but a secondary effect in translation efficiency (Fiszbein et al., 2019), we expect an even stronger effect at the protein level. Our mechanism of increased mRNA and protein expression through splicing could potentially explain the increase in SMN2 endogenous levels in the original study targeting exon 7 inclusion with ASOs (Hua et al., 2007).

The catalogue of human EMATS genes that we provided here serves as an example of the thousands of genes that are sensitive to this approach. Since the small molecule or ASO should be able to increase inclusion of an alternative exon correctly positioned, the activation of gene expression through splicing is limited to SE in EMATS genes. However, a similar strategy used to inhibit gene expression through splicing would also work for constitutive exons in which splicing can be downregulated. The usage of constitutive exons to inhibit gene expression through splicing would expand the number of genes sensitive to a similar approach by several folds. Altogether, our findings provide evidence for the development of the first therapeutic strategy to increase gene expression through splicing by EMATS, as well as a list of ∼1500 genes in which this approach presents the strongest effects.

## Acknowledgments

We thank Christopher B. Burge for helpful discussions, ideas, and comments, as well as members of the Fiszbein lab for feedback on the manuscript. This work was supported by university funds and grants to A.F from the NIH (1R35GM147254-01) and the Richard and Susan Smith Family Foundation.

## Author Contributions

M.U.A. performed all experiments and analyses in Figures 4, 5, and 6. S.T.M. developed the code to identify EMATS genes and performed all computational analyses in Figures 1, 2 and 3 with help from Z.S. and R.R.. A.F. designed the study, wrote the manuscript, and supervised the entire work.

