## Supplemental material for "Splicing-dependent transcriptional activation"

Supplemental figures

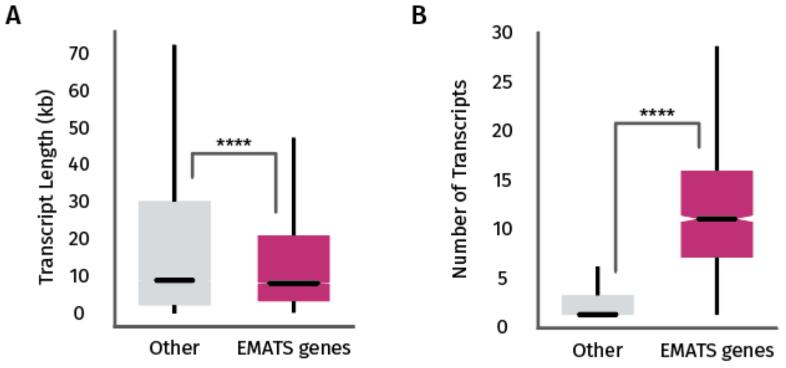


**Figure S1**

**A.** distribution of total transcript length for isoforms of genes with EMATS structure and other genes. **B.** distribution of the number of isoforms per gene for genes with EMATS structure and other genes.

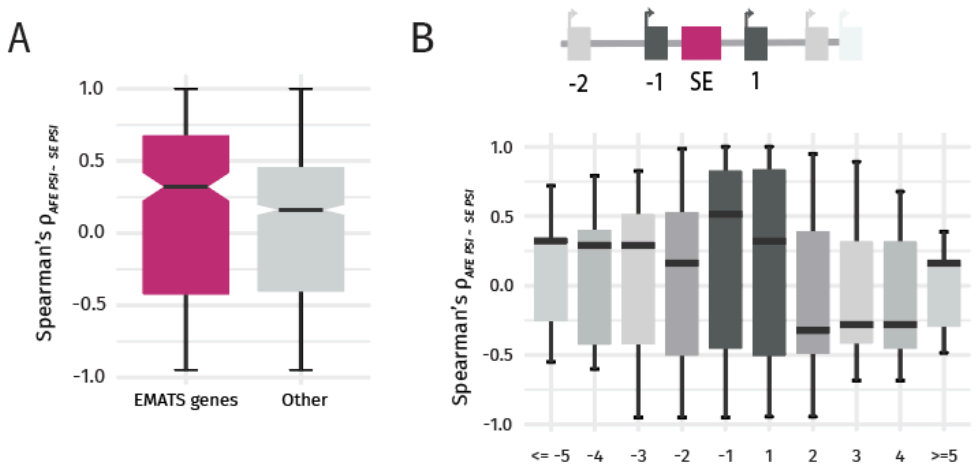


**Figure S2**

Distribution of the spearman correlation between changes in gene expression and changes in SE PSI during SARs-Covid2 infection for non-EMATS genes compared to EMATS genes (**A**) binned by the position of the promoter relative to the position of the SE located in an EMATS structured (**B**). Promoters are numbered with increasing positive values downstream of the SE and with decreasing negative values upstream of the SE.

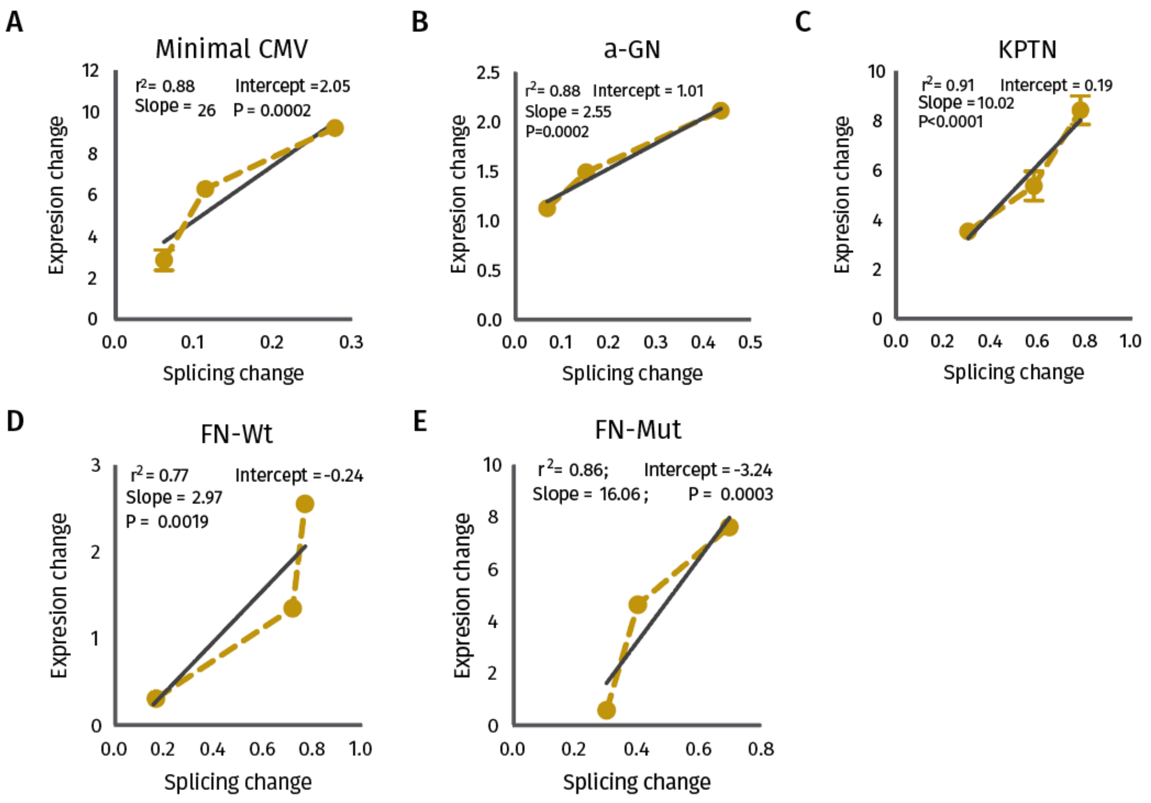


Figure S3

Correlation between splicing changes and gene expression after Risdiplam treatment with three different concentrations driven by a minimal CMV promoter (**A**), an alfa-globin (a-GN) promoter (**B**), a KTPN promoter (**C**), the fibronectin wilt type (FN-Wt) promoter (**D**), and the fibronectin mutated (FN-Mut) promoter (**E**).

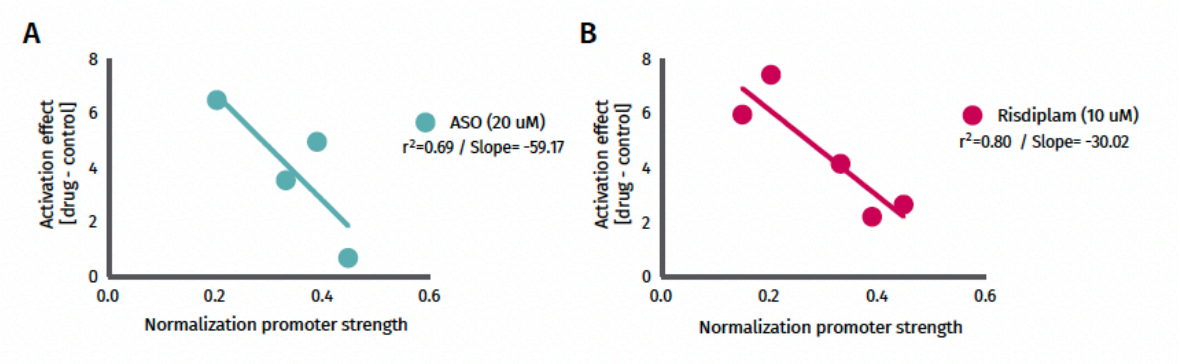


Figure S4

Linear regression of gene expression changes in SMN2 splicing reporter following ASO treatment (**A**) and Risdiplam treatment (**B**) for stable lines under the regulation of the four different human promoters tested and promoter strength measured by basal RT-qPCR in control conditions.
